# Regulation of miR-219 and its target RalGPS dictates acute synaptic plasticity at the *Drosophila* neuromuscular junction

**DOI:** 10.1101/2025.11.06.686960

**Authors:** Kumar Aavula, Hansine Heggeness, Takakazu Yokokura, David Van Vactor

## Abstract

RNA regulatory mechanisms mediate adaptive responses requiring rapid gene expression changes. Examining microRNA (miRNA)-mediated control of structural synaptic plasticity, we identified miR-219 and six other conserved miRNAs regulating activity-induced morphogenesis at *Drosophila* glutamatergic synapses. Among miRNA targets downregulated by acute stimulation, the Ral-activating factor RalGPS is necessary for presynaptic morphogenesis and sufficient to block plasticity. Stimulation elevates miR-219 loading into Argonaute complexes without altering mature miR-219 levels, revealing a post-transcriptional regulatory mechanism. We conclude that activity-induced miRNA loading modulates Ral signaling via RalGPS to enable presynaptic remodeling and growth, establishing a novel link between neuronal activity and synaptic structural plasticity.

## Introduction

Long-term information storage within neural circuits relies on morphological changes in synaptic architecture initiated by transient, activity-dependent alterations in gene expression (Kandel 2001). Among mechanisms controlling neuronal gene expression, non-coding miRNAs offer temporal and spatial versatility in responses to synaptic signaling (McNeill and Van Vactor 2012; Soutschek and Schratt 2023). The nervous system exhibits the richest landscape of miRNA expression across animal species (Dubes et al. 2019; Hu and Li 2017), and transcriptome profiling has revealed that neuronal activity regulates miRNA expression (Nesler et al. 2013; Portugal et al. 2025; Pichardo-Casas et al. 2012; Spronsen et al. 2013). Leveraging such profiling data and cell-culture models, substantial progress has been made in understanding miRNA function in postsynaptic structures such as dendrites and dendritic spines (Soutschek and Schratt 2023). However, despite recent insights into miRNA control of synaptic homeostasis (Rajman et al. 2016; Srinivasan et al. 2021; Dubes et al. 2019; Fiore et al. 2014; Colameo et al. 2025), the roles of miRNAs in presynaptic plasticity—particularly structural remodeling—remain poorly understood (Lu et al. 2014; McNeill et al. 2020; Nesler et al. 2016; Siegert et al. 2017).

Genetic models such as *C. elegans* and *Drosophila* provide powerful systems for surveying miRNA functions in vivo, offering repositories of mutations and transgenic tools for conditional perturbation (e.g. Fulga et al. 2015; Bejarano et al. 2012). Early studies uncovered the first miRNA regulators of synaptic form and function, highlighting postsynaptic roles for *C. elegans* miR-1 and *Drosophila* miR-8 (Loya et al. 2009; Simon et al. 2008). Subsequent profiling in *Drosophila* revealed that neuronal activity controls mature levels of several miRNAs (Nesler et al. 2013), including miR-8, which is essential for activity-dependent remodeling of glutamatergic synapses at the larval neuromuscular junction (NMJ), as well as miR-289 (Nesler et al. 2016). However, unbiased anatomical screening identified over a dozen additional miRNAs required for NMJ development (McNeill et al. 2020), raising the question of whether many miRNAs contribute to synaptic plasticity independent of activity-dependent changes in their expression.

*Drosophila* NMJ morphogenesis is influenced by neural activity during larval development (Van Vactor and Sigrist 2017; Menon et al. 2013; Koh et al. 2000). Using neuron-specific expression of transgenic competitive “sponge” inhibitors (miR-SP) that identified synaptic phenotypes in our prior screen (McNeill et al. 2020), we tested whether these miRNAs regulate bouton initiation in response to elevated activity. Our current results indicate that more than 70% of miRNAs with developmental phenotypes also control acute activity-stimulated growth. Focusing on one highly conserved candidate, we found that miR-219 downregulates the Ral GTPase activator RalGPS. Both elevation and elimination of RalGPS prevent activity-induced synaptic remodeling and normal presynaptic terminal formation, suggesting that Ral activity must be precisely tuned in the presynaptic compartment.

## Results and Discussion

### Activity-induced Synaptic Growth at the Larval NMJ is Extensively Regulated by miRNA

To determine if miRNAs identified in our prior NMJ screen (McNeill et al. 2020) control some presynaptic aspect of activity-induced remodeling, we expressed miR-SP lines in motor neurons and then subjected larvae to a well-established spaced depolarization protocol (Ataman et al. 2008). This method, consisting of multiple cycles of high K⁺ stimulation followed by brief rest periods, induces the formation of immature, nascent synaptic boutons, also known as ghost boutons. Ghost boutons are defined by their lack of postsynaptic density cytomatrix (Fig. 1C), and serve as a reliable feature of acute synaptic structural plasticity (Ataman et al. 2008; Piccioli and Littleton 2014). Analysis of nine miRNAs validated in our prior study (McNeill et al., 2020) revealed that knockdown of miR-13a, miR-14, miR-34, miR-219, miR-277 and miR-973 significantly impaired new bouton formation in response to acute stimulation without altering the baseline number of nascent boutons, whereas inhibition of miR-1014 led to a significant increase in activity-induced nascent boutons (Fig. 1A; Fig. S1G shows the percentage change from control). Inhibiting miR-316 failed to alter presynaptic activity-induced remodeling in this acute assay (Fig. 1A). These data indicate that 78% of miRNAs detected in our previous screen regulate activity-dependent synaptic remodeling.

**Figure 1.**
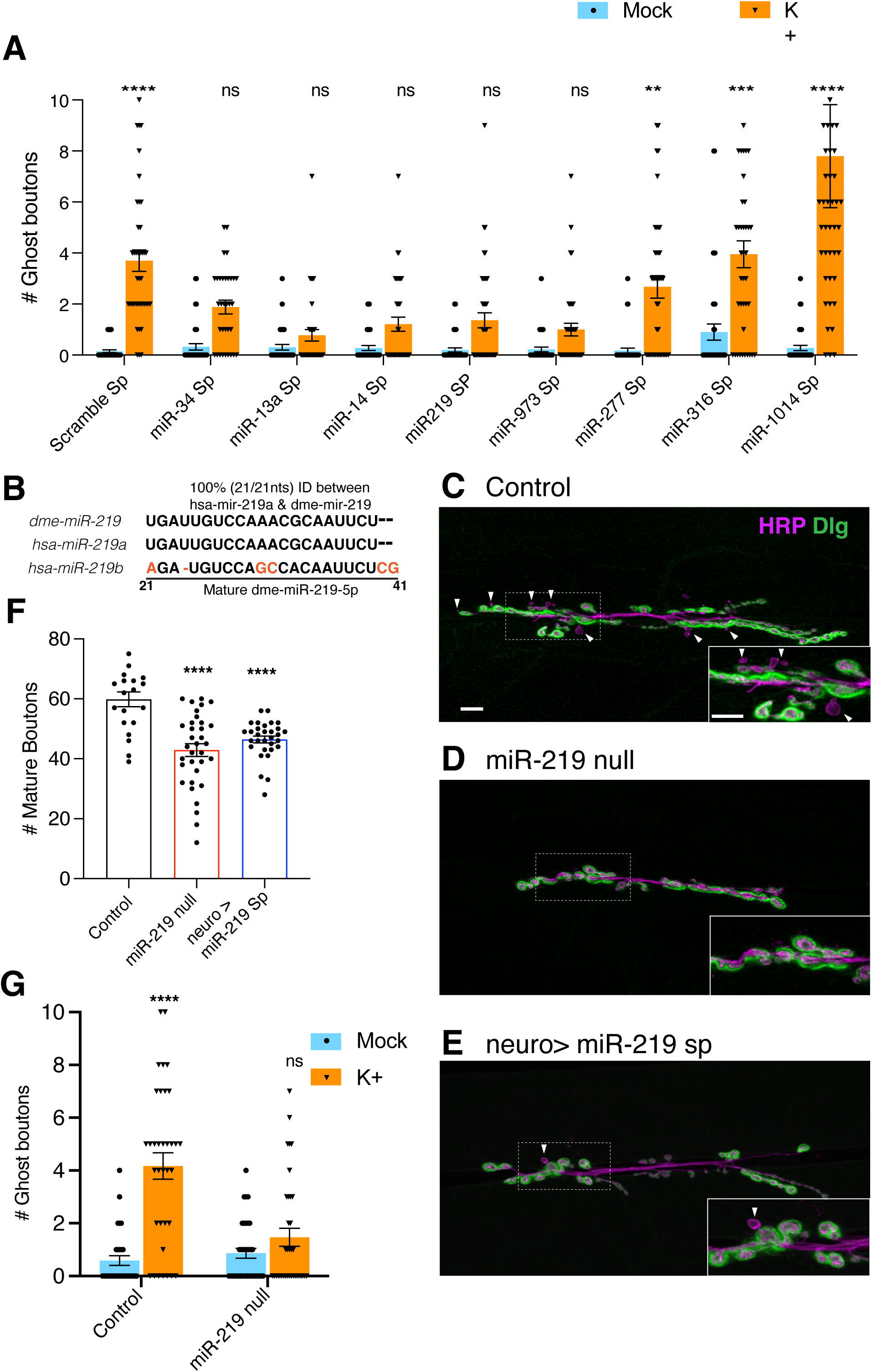
miR-219 is essential for synaptic plasticity at neuromuscular junction. **A.** Screen to identify miRNAs that are required for acute synaptic structural plasticity in motor neurons using high K+ (90mM) induced membrane depolarization assay. Mean ± SEM, two-way ANOVA , Sidak’s multiple comparisons. **B.** Sequence alignment showing 100% identity between *Drosophila miR-219-5p* and human *miR-219a-5p*. **C–E.** Confocal images of NMJ arbors on muscles 6/7 after stimulation with high K⁺, stained for presynaptic membrane (HRP, magenta) and postsynaptic cytomatrix (Dlg, green). Arrowheads indicate ghost boutons **C.** Control, **D.** miR-219 null mutant, **E.** Expression of *miR-219-5p* sponge (SP) in motor neurons (*OK371-GAL4/UAS-miR-219SP; UAS-miR-219SP/+*). Scale bars: 10 μm. **F.** Quantification of mature boutons. Loss of *miR-219* in motor neurons significantly reduces number of mature boutons. Mean ± SEM, One-way ANOVA and Tukey’s multiple comparisons. **G.** Quantification of ghost boutons after high K⁺ stimulation. Loss of miR-219 fails to induce bouton addition after stimulation. Mean ± SEM, Two-way ANOVA and Sidak’s multiple comparisons. n= 40 NMJ from 10 animals. Statistical significance: ****p ≤ 0.0001, ***p ≤ 0.001, **p ≤ 0.01, *p ≤ 0.05, ns = not significant (p ≥ 0.05).

Although some analysis has been performed for miR-34 at the NMJ (McNeill et al. 2020) and previously for miR-8 and miR-289(Nesler et al. 2013, 2016), little is known about the presynaptic functions of most miRNAs identified in our targeted screen. Among the miRNAs required in motor neurons for the formation of nascent boutons in response to high K^+^, we were particularly interested in miR-219 due to an exceptionally high degree of conservation between fly and human (mature dme-miR-219 is 100% identical to hsa-miR-219a; see Fig 1B). When we compared neuronal-specific inhibition of miR-219 to a complete null mutation (KO) eliminating all miR-219 expression, we found the phenotypes to be both qualitatively and quantitatively comparable for mature NMJ structure (Fig 1C-F), as well as for activity-induced bouton addition (compare Fig 1A to Fig 1G). This confirmed that miR-219 is required for normal synapse growth and that a significant component of this function is cell-autonomous to presynaptic neurons. Interestingly, studies in mammalian systems suggest that miR-219 is a vital regulator of neuronal and glial cell biology, and it has also been implicated in schizophrenia, depression, and multiple sclerosis(Dugas et al. 2010; Kocerha et al. 2009). Moreover, miR-219 has been found to target multiple neuronal mRNAs, including NMDA-class glutamate receptors and the Calcium-Calmodulin-dependent protein kinase II [CaMKII](Kocerha et al. 2009; Pan et al. 2014; Wang et al. 2018) in mammals, as well as the Fragile-X ortholog [dFMR1] and Tau in *Drosophila* (Wang et al. 2019; Santa-Maria et al. 2015). However, in contrast to *miR-219* itself, none of the published target mRNAs for *miR-219* contain miRNA response element(s) (MREs) that are conserved from fly to man.

### The Guanine-Nucleotide Exchange Factor RalGPS is a potential target of miR-219

To find downstream targets of miR-219 that mediate activity-induced synaptic plasticity, we first performed RNA sequencing on total RNA extracted from larval pelts of *miR-219* null mutants compared to genetically matched wild type control samples. Biological replicates of these genotypes displayed clear segregation by hierarchical clustering (Fig S2A). Differential expression analysis revealed over 1,000 genes were significantly upregulated in the absence of miR-219. Gene Ontology (GO) enrichment analysis indicated that 70% of the enriched GO categories are associated with synaptic signaling pathways, particularly those related to glutamatergic transmission and receptor pathways known to be relevant to neuronal plasticity (Fig. 2A). However, only a subset of these genes were potential direct targets for miR-219; specifically, 28 of the upregulated transcripts were predicted by TargetScanFly to contain miR-219 MRE sequences (Fig. 2B and Fig. S2B). We then asked if miR-219 MREs found in upregulated transcripts were also conserved in the human orthologs of the predicted target genes. Only four of 28 candidates displayed MRE conservation in their mammalian cognates: NaCP60E (a voltage-gated sodium channel), Elk (a voltage-gated potassium channel), CG31523, and CG5522 (a guanine nucleotide exchange factor [GEF], RalGPS) (Fig. 2B, Fig. S2B).

**Figure 2.**
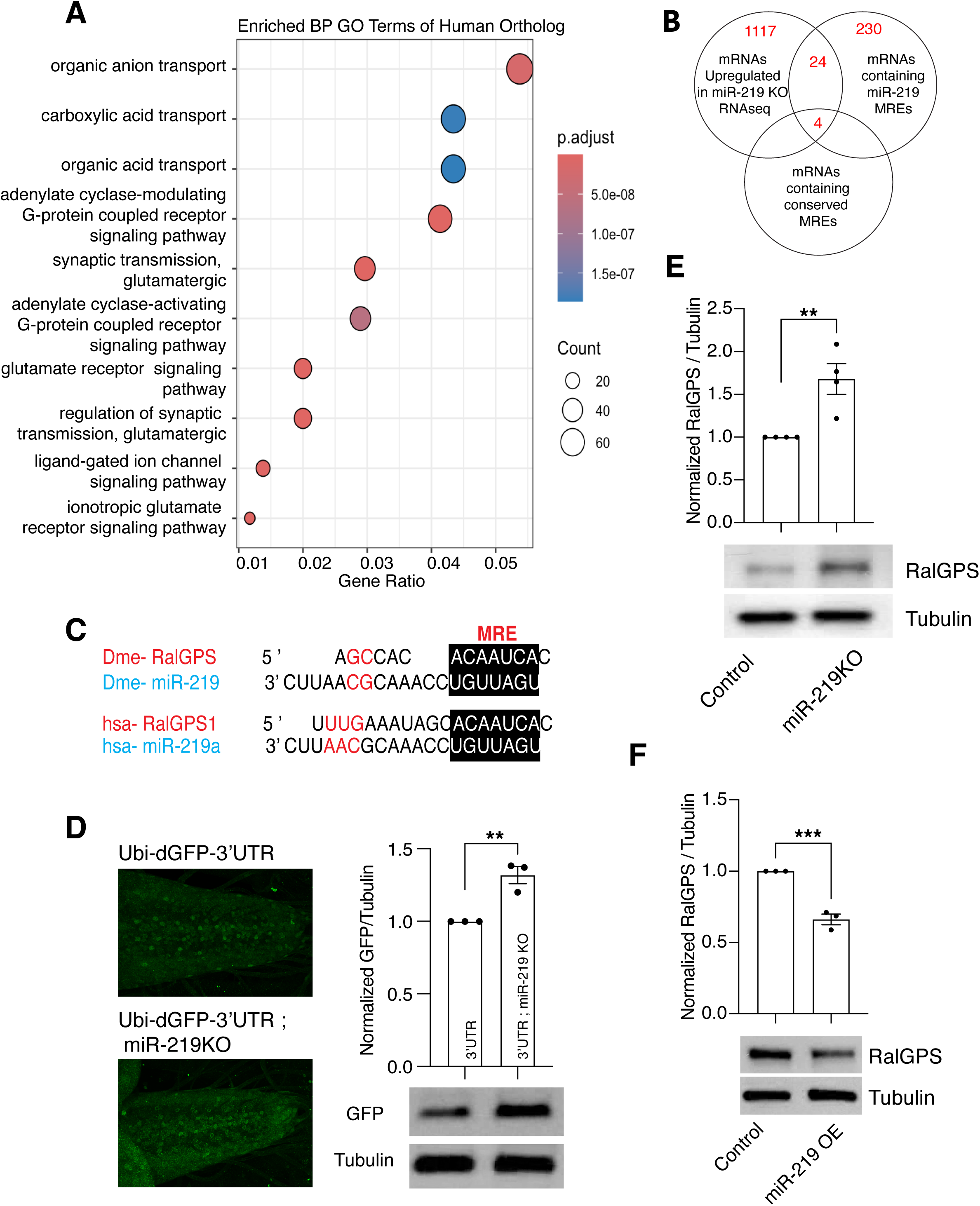
RalGPS is a functional target of miR-219. **A.** Functional enrichment analysis of biological process terms associated with human orthologs of *Drosophila* genes upregulated in miR-219 null RNA-seq. **B.** A Venn diagram depicting the overlapping criteria applied to refine and identify potential targets of miR-219. **C .** Alignment showing conserved miR-219 seed and MRE elements in *Drosophila* RalGPS and human RalGPS1. **D.** Analysis of the larval ventral nerve cord ubiquitously expressing GFP linked to the RalGPS 3′UTR in both wild type and miR-219 null backgrounds, accompanied by Western blot validation and quantification of GFP expression. **E-F.** Western blot analysis of expression of RalGPS (RalGPS-3xHA) in wildtype, *M190sl* null (**E**) and overexpression of miR-219 in motor neurons (**F**). Mean ± SEM, Student’s t-test. Statistical significance: ***p ≤ 0.001, **p ≤ 0.01.

Because GO enrichment analysis suggested significant involvement of genes acting in synaptic signaling pathways, we selected RalGPS for further analysis. RalGPS is a highly conserved GEF known for its Ras-independent activation of the Ral GTPase (Rebhun et al. 2000; Peng et al. 2011), though little is known about its synaptic function; moreover, the *Drosophila* ortholog (CG5522) has not been studied in the nervous system. In the 3’ untranslated region (UTR) of *Drosophila* and human RalGPS/RalGPS1 mRNAs, a single MRE for *miR-219/miR-219a* is present with a perfect match 7-mer-A1 seed sequence (Fig 2C). We introduced the *Drosophila* RalGPS 3’UTR downstream of a short-lived GFP cDNA (dGFP; (He et al. 2019)) under control of a ubiquitin promotor (Ubi; pUWR, DGRC#1281) to assess the miR-219-dependence of this UTR sequence for reporter expression in vivo (see Methods). When the Ubi-dGFP-RalGPS 3’UTR reporter was introduced into a *miR-219 KO* null mutant background, the level of GFP expression increased significantly in the larval central nervous system ([CNS] consisting of ventral nerve cord [VNC] and brain) as assessed by fluorescence microscopy or quantification of Western blots with anti-GFP antibody, compared to the same reporter in a matched wild type control background (Fig 2D). This suggested that RalGPS can serve as a direct target of miR-219 downregulation. To confirm that RalGPS expression is repressed by miR-219 at the protein level, we used CRISPR-Cas9 to epitope tag the endogenous RalGPS at the C-terminus; Western blot analysis of larval brains revealed CNS expression (see Methods; Fig. S2C). We then compared the expression of the tagged RalGPS protein in genetic backgrounds wild type or null for *miR-219*. As predicted by our UTR analysis, RalGPS protein was significantly increased in larval pelts from *miR-219 KO* (Fig 2E) while it was decreased when *miR-219* was overexpressed chronically in motor neurons (Fig 2F). Together, our observations suggest that RalGPS serves as a target of *miR-219*.

### miR-219 and RalGPS are Activity-Dependent

The requirement of miR-219 in motor neurons for bouton addition induced by acute stimulation (Fig. 1) raised the question of whether the levels of this miRNA might respond to neural activity. We used TAQMAN qPCR on RNA extracted from the larval pelt after stimulation with K^+^ spaced depolarization to quantify mature miR-219 miRNA compared to mock-stimulated controls. Yet, we found no significant change in miR-219 levels after stimulation (Fig. 3A, center bar). Because miR-34 is required for NMJ growth and activity-dependent bouton addition, whereas let-7 is not required for NMJ growth (McNeill et al. 2020), we also measured mature miRNA levels for these, but neither showed any significant change after stimulation (Fig 3A); these data were consistent with activity-dependent profiling performed previously by microarray (Nesler et al. 2013), suggesting that there is no major increase in processing of let-7, miR-34 and miR-219 pre-miRNA species in the acutely stimulated larval pelt.

**Figure 3.**
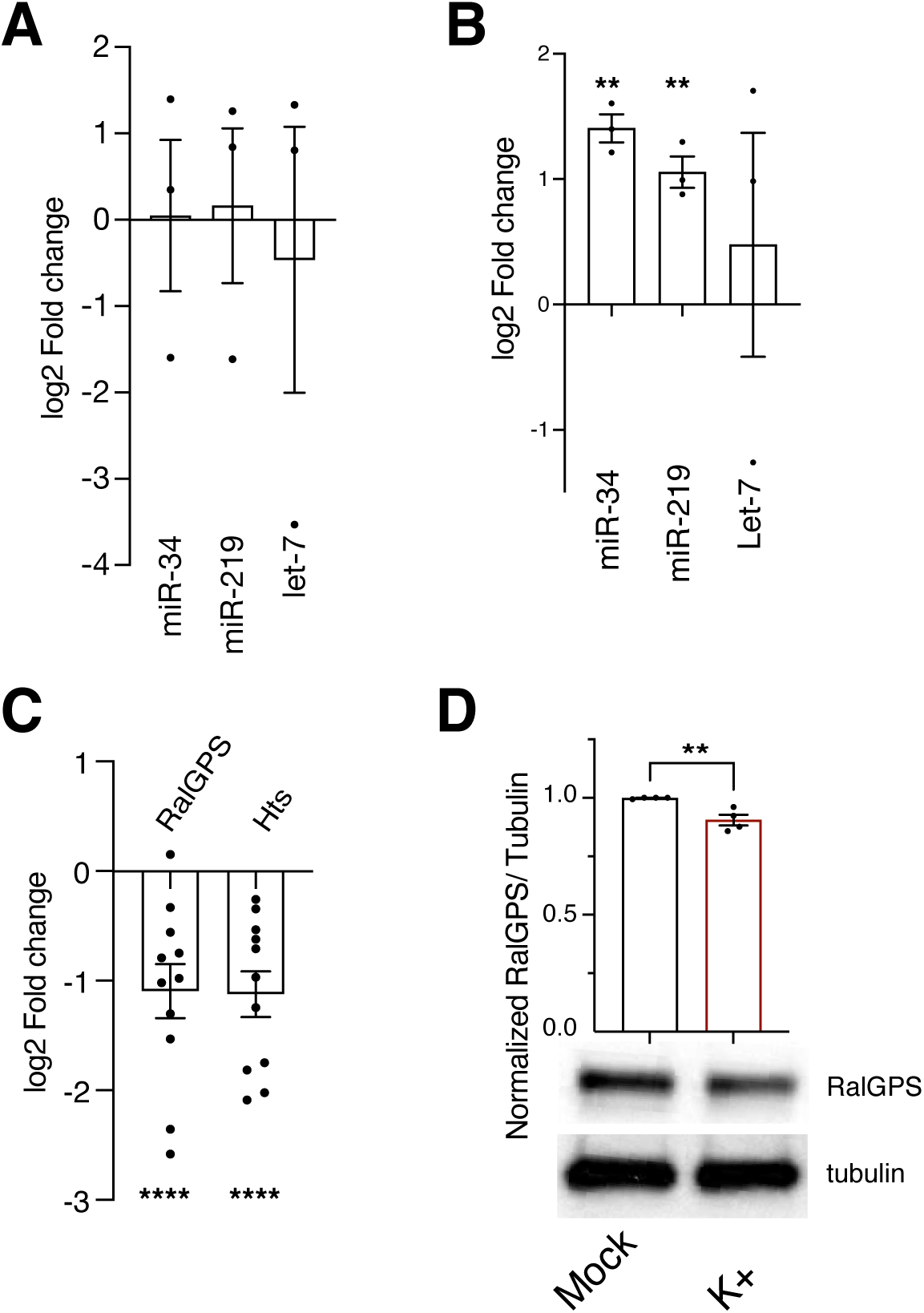
Activity-dependent regulation of miR-219 and RalGPS. **A.** TaqMan qPCR showing total miRNA levels in RNA from K⁺-treated larval pelts. **B.** TaqMan qPCR of miRNAs co-immunoprecipitated with Ago1, showing increased miRNA loading into the Argonaute complex following K⁺ stimulation. Mean ± SEM, Welch’s t-test. **C.** qPCR showing downregulation of *RalGPS* mRNA after K⁺ stimulation; *Hts* is used as positive control. Mean ± SEM, One-way ANOVA and Dunnett’s multiple comparisons. **D.** Western blot showing reduced RalGPS protein levels upon K⁺ stimulation. Mean ± SEM, Student’s t-test. Statistical significance: ****p ≤ 0.0001***p ≤ 0.001, **p ≤ 0.01, *p ≤ 0.05, ns = not significant (p ≥ 0.05).

Because active miRNA species must load into the miRNA-specific RNA-Induced Silencing Complex (miRISC) in order to pair and regulate target RNAs (Dexheimer and Cochella 2020; Jungers and Djuranovic 2022), and miRISC activity has been shown to respond to activity in prior studies(Nawalpuri et al. 2020; Srinivasan et al. 2021), we wondered if miR-219 loading might change after acute stimulation. We obtained and tested several antibodies directed against the *Drosophila* miRNA-selective RISC component Argonaut-1 (Ago1), and found two independent antibodies capable of efficient and specific Ago1 immunoprecipitation (IP) from larval pelt tissue (Fig. S3A). Using these antibodies to pull down miRISC from pelt lysates, followed by TAQMAN qPCR to detect mature miRISC-bound miRNA species, we compared the loading of miR-219 in acutely stimulated versus mock-stimulated larvae. Here we found that there was a significant increase in miR-219 loading, with a nearly two-fold change (Fig. 3B, center bar; p<0.01). Interestingly, when we quantified mature miR-34 and let-7 from the same Ago1/miRISC IPs, we saw a comparable increase in miR-34 loading (p<0.01), but no significant change in let-7 (Fig. 3B). Our observations suggest that stimulation increases the activity of miR-219 in our system.

The finding that RalGPS expression is dependent on miR-219 (Fig. 2) and that miR-219 loading is activity-dependent led to a prediction that RalGPS expression might be responsive to neural activity. We first tested this notion at the RNA level, using rt-qPCR to quantify RalGPS transcripts in acutely-stimulated versus mock-stimulated larval pelt samples. Consistent with our other data, we found that RalGPS mRNA levels decrease by approximately the same degree of change in stimulated pelts as the increase in miR-219 loading (compare log2-fold change in Fig. 3B to Fig. 3C). Because miR-34 also showed an activity-dependent increase in miRISC loading, we quantified the levels of the known miR-34 target gene Adducin/Hts (McNeill et al. 2020); Hts transcripts decreased upon acute stimulation by roughly the same amplitude as the reduction in RalGPS (Fig. 3C), consistent with the activity-dependent increase in miR-34 observed in our Ago1 IPs. We next used our tagged endogenous RalGPS strain to ask if we could detect any change in RalGPS protein level in the acute stimulation preparation. Western blots comparing mock-stimulated to K+-stimulated tissue did show a highly significant decrease in RalGPS (Fig. 3D), albeit at an effect size less than the change in RalGPS mRNA. Therefore, RalGPS is an activity-regulated gene.

### RalGPS Plays an Essential Role in Synapse Morphogenesis and Plasticity

If RalGPS is a miR-219 target gene whose down regulation is important for activity-induced synapse growth, then we reasoned that elevation of RalGPS in otherwise normal motor neurons might interfere with bouton addition in response to stimulation. Using a UAS-dme-RalGPS cDNA transgene combined with a motor neuron-specific Gal4 driver, we found that elevated RalGPS did not significantly alter the baseline level of ghost boutons in late larval NMJs. However, when we compared mock-stimulated and K+-stimulated animals, we found that increasing RalGPS expression reduced ghost bouton addition over two-fold compared to a Gal4 alone control genotype (Fig 4A). This suggests that RalGPS is sufficient to inhibit synaptic remodeling.

**Figure 4.**
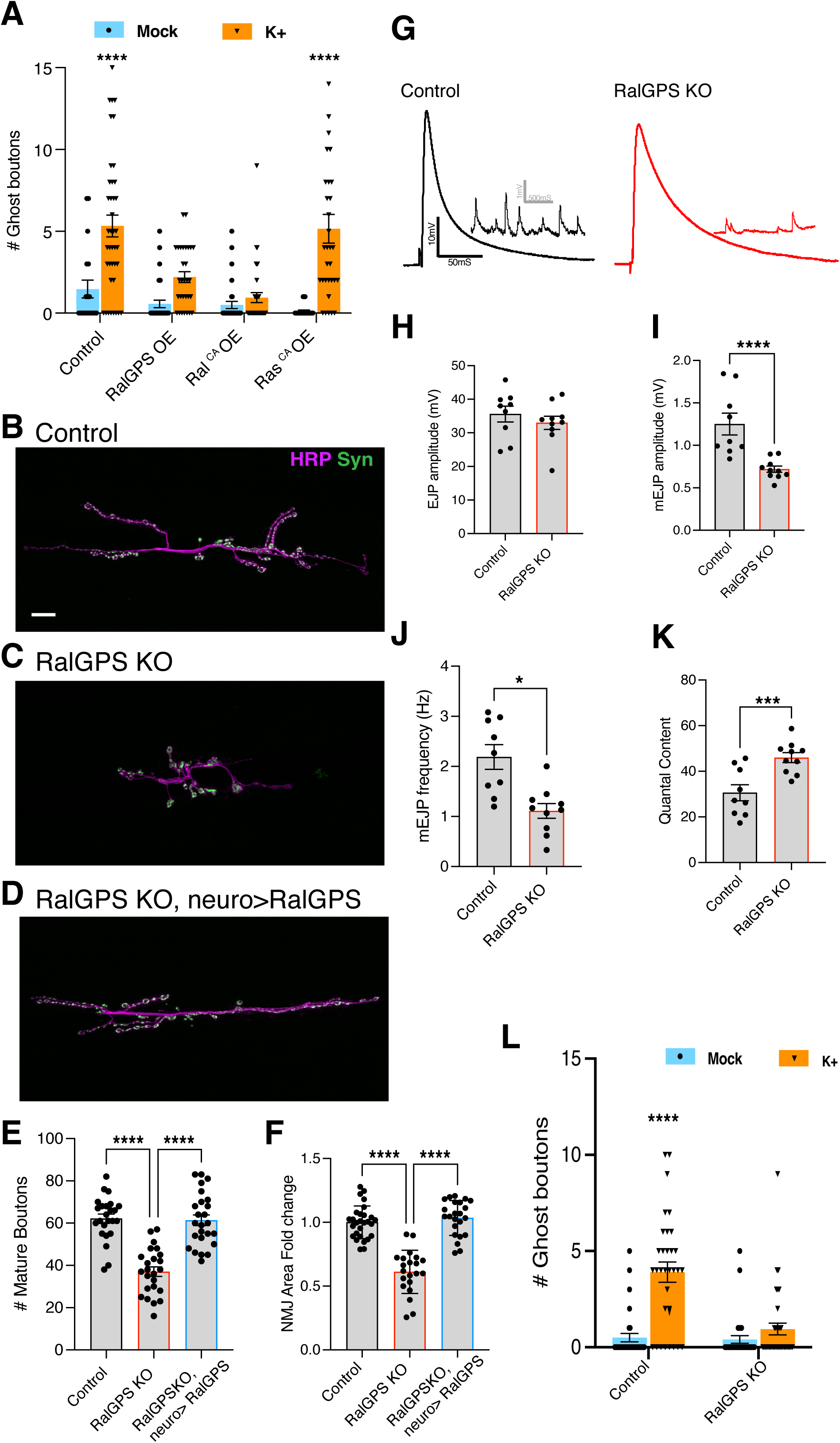
RalGPS is essential for synapse development and plasticity. **A.** Quantification of ghost boutons after K⁺ stimulation. Overexpression of *RalGPS* or constitutively active RalA in motor neurons inhibits induction of ghost boutons. Mean ± SEM, two-way ANOVA , Sidak’s multiple comparisons**. B–D.** Confocal images of NMJs (muscles 6 /7), stained for presynaptic membrane (HRP, magenta) and synaptic vesicles (Syn, green) **B.** Control NMJ with well-organized boutons **C.** Loss of *RalGPS* results in reduced bouton number **D.** Motor neuron expression of *RalGPS* rescued bouton number in *RalGPS* null. Scale bars: 10 μm. **E.** Quantification of number of mature boutons and **F.** Fold change in NMJ area. Mean ± SEM, One-way ANOVA and Tukey’s multiple comparisons. **G.** Representative traces of evoked (EJP) and miniature (mEJP) junctional potentials. **H.** Quantification of EJP amplitudes shows no significant difference in *RalGPS* mutants compared with controls. **I.** mEJP amplitudes are significantly reduced in *RalGPS* mutants. **J.** mEJP frequency is significantly decreased, whereas **K.** quantal content is significantly increased in *RalGPS* mutants. Mean ± SEM, Student’s t-test. **L**. Quantification of ghost boutons after K⁺ stimulation in *RalGPS* null shows that Loss of *RalGPS* fails to induce ghost boutons. Mean ± SEM, two-way ANOVA , Sidak’s multiple comparisons. Statistical significance: ****p ≤ 0.0001***p ≤ 0.001, **p ≤ 0.01, *p ≤ 0.05, ns = not significant (p ≥ 0.05).

RalGPS encodes a conserved GEF specific to Ral-family GTPases (Rebhun et al. 2000; Peng et al. 2011)(protein sequence alignment between fly and human RalGPS is shown in Fig. S4A, highlighting the predicted binding site for Ral in Fig. S4B). Thus, given our discovery that elevated RalGPS is sufficient to block synapse remodeling, we predicted that activation of Ral should also be sufficient to prevent bouton addition in response to acute stimulation. When we used motor neuron-specific Gal4 to express a constitutively active Ral transgene (UAS-Ral^CA^; (Teodoro et al. 2013)), we saw complete suppression of bouton addition in response to spaced depolarization (Fig. 4A). A key defining feature of the RalGPS family of GEFs is absence of the Ras-Activation (RA) domain that links other Ral GEFs (e.g. RalGDS) to active Ras(Peng et al. 2011). This unique independence of RalGPS from Ras (compared to all other families of Ral GEF), suggested that bouton addition in response to acute stimulation should be Ras independent if the key mechanism was governed by miR-219 downregulation of RalGPS leading to a decrease in Ral activity in motor neurons. To investigate the logic of the mechanism, we expressed a constitutively active Ras (UAS-Ras^CA^; (Coleman-Gosser et al. 2023)) in motor neurons and then subjected these animals to mock or acute K^+^ stimulation. Interestingly, elevation of Ras activity did not block bouton addition upon high K+ treatment (Fig. 4A) despite the fact that it plays a role in NMJ development (Koh et al., 2002), suggesting that Ral activity is under the control of RalGPS and miR-219, but not Ras, for new bouton morphogenesis in motor neurons that have been acutely stimulated.

The ability of increased RalGPS and Ral activity to block morphological plasticity raised questions of (a) whether RalGPS might act simply as a brake or lock to prevent remodeling under baseline conditions, and, (b) what this signaling pathway is required for under baseline conditions? As there were no existing null mutations in *Drosophila* RalGPS, we used CRISPR-Cas9 to generate a deletion of the coding sequence for the gene with guide RNAs flanking the coding exons (see Fig. S4C). The null animals were viable with no striking external developmental abnormalities that we could detect at adult, pupal or larval stages (see Methods). However, at the late third instar, immunostaining of larval pelts revealed a significant reduction in overall NMJ area and the number of mature boutons in *RalGPS KO* nulls (compare Fig. 4B to Fig. 4C; quantified in Fig. 4E-F). Both the reduction in bouton number and the decrease in NMJ arbor area were effectively rescued by add-back of a wild type RalGPS cDNA specifically in motor neurons (Fig. 4D-F), confirming a presynaptic requirement for the gene.

Given the striking morphological NMJ phenotype in the *RalGPS KO*, we next asked if these null synapses are capable of neurotransmission. Electrophysiological recordings demonstrated that evoked excitatory junctional potentials (EJPs) are not significantly reduced in amplitude in the *RalGPS KO* homozygotes compared to controls (Fig. 4G shows representative traces; quantified in Fig. 4H); although the kinetics of recovery to resting potential is slower in *RalGPS* mutants (see Methods). Interestingly, recordings of spontaneous release events (miniature mEJPs) showed a significant reduction in both amplitude (Fig. 4I) and frequency (Fig. 4J), suggesting that the increase in quantal content in *RalGPS KO* (Fig. 4K) probably represents a RalGPS-independent adaptive compensation maintaining relatively normal evoked output (Fig. 4H). Thus, as has been observed in many other mutant backgrounds(Albin and Davis 2004; Mushtaq et al. 2022), despite substantial defects in NMJ structure, these synapses are functionally robust. However, the striking decrease in boutons raised the question of whether the *RalGPS KO* mutants are solely defective in baseline developmental bouton addition or whether the NMJ growth defect seen in the null might also reflect a failure in NMJ plasticity. When we subjected the *RalGPS KO* larvae to the spaced depolarization assay, we found a near-complete loss of acute activity-stimulated bouton addition (Fig. 4L). Therefore, chronic elevation or loss of RalGPS, and presumably Ral GTPase activity, are both deleterious to activity-dependent NMJ plasticity.

Early studies of Ras function at the *Drosophila* NMJ indicated that the mitogen-activated protein kinase (MAPK)-selective downstream output of Ras mediates its major long-term impact on matching developmental synapse growth with muscle growth (Koh et al. 2002). Although contributions of other Ras effector pathways were less than that MAPK, the study showed that Ral also promotes NMJ growth, possibly via activation of phospholipase D (PLD)(Koh et al., 2002); however, the acute K+ stimulation protocol was developed later (Ataman et al., 2008). Our analysis suggests that rapid activity-induced bouton addition depends on regulation of Ral but not Ras, implying that these GTPases are not always part of a requisite cascade. In the acute context, we find that RalGPS is a key factor controlling NMJ morphogenesis. It remains unclear what upstream signals control Ras during NMJ development, but our data suggest that miR-219, and potentially other factors, regulate presynaptic Ral. Interestingly, the conserved Plexstrin-Homology (PH) and proline rich SH3-binding motifs in RalGPS may implicate phosphoinositides and SH3-domain proteins as modulators of RalGPS activity and/or localization (Rebhun et al. 2000). This leaves open the question of whether a convergence of signals collaborates with miR-219 to dynamically modulate Ral activation by RalGPS in motor neurons.

## Acknowledgements

We are grateful to our colleagues Drs. Spyros Artavanis-Tsakonas and Danesh Moazed for helpful feedback on this manuscript. We thank Dr. Pushpa Verma and Mr. Niket Govaram for pilot experiments with tagged Ago transgenics to test the IP protocol. We are grateful to Dr. Howard Lipshitz and colleagues for their anti-Ago1 fab fragment clones, and to Dr. Mikiko Siomi for generous gifts of a published anti-dme-Ago1 monoclonal antibody that we used in optimizing our IP protocol. We thank the CITE imaging core and its staff at Harvard Medical School for access to imaging instrumentation. Much of our work was dependent on government-supported resources such as Flybase (https://flybase.org/), the Bloomington *Drosophila* Stock Center (https://bdsc.indiana.edu), and the Developmental Studies Hybridoma Bank (https://dshb.biology.uiowa.edu/). This study was supported by a grant from NINDS (NS135403) and internal resources from Harvard Medical School.

## Author contributions

KA and DVV conceived of the project. KA designed and performed all aspects of the project and analysis of the data. HH assisted with imaging experiments and data analysis, and also performed all of the sequence analysis and informatics; TY contributed to construction of reporter and other expression constructs; KA and DVV wrote the manuscript.

## Methods

### Fly stocks

Flies were raised on standard fly food at room temperature. All genetic crosses were performed at 25°C. w^1118^ or parental stock crossed with w^1118^ was used as controls for mutants and crosses respectively. The miRNA and scramble sponge stocks were generated by Van Vactor lab(Fulga et al. 2015) . *miR-219* null (BDSC#58900), UAS-miR-219 (BDSC#59890), Ok371-Gal4 (BDSC#26160), UAS-Ral^CA^ (UAS-Rala^G20V^, BDSC#81049), UAS-Ras^CA^ (UAS-Ras85D^V12^, BDSC# 64196), were obtained from Bloomington stock center (BDSC, Bloomington, IN, USA).

### High K⁺ Depolarization and Ghost Bouton Analysis

High K⁺-induced depolarization was performed using HL3 buffer containing 90 mM K⁺. Wandering third instar larvae were dissected and subjected to five cycles of stimulation, each consisting of 5 minutes in 90 mM K⁺ followed by a 15-minute rest. Following stimulation, larval fillets were fixed in Bouin’s solution and immunostained for presynaptic membrane (HRP) and postsynaptic cytomatrix (Dlg). Ghost boutons were imaged on muscles 6 and 7 of abdominal segments A3 and A4 using a Nikon A1R point-scanning confocal microscope and a Nikon 90i fluorescence microscope. Quantification was performed based on standard criteria for ghost bouton morphology (Ataman et al. 2008) .

### CRISPR-Mediated Generation of RalGPS Deletion and HA-Tagged Transgenes

CRISPR target sites for the RalGPS locus were identified using the FlyCRISPR Target Finder tool (flycrispr.org/target-finder). Two guide RNAs (gRNAs) flanking the RalGPS gene were selected to generate a precise deletion without disrupting adjacent genes: 5′ gRNA: GCAAGCCGAAACGAAGCTTC, 3′ gRNA: GTCCGTCCGGAAGCGATAGA.

gRNAs and donor plasmids were designed and cloned following established protocols (Gratz et al. 2014) , using the following primers:

gRNA cloning oligos:

5’ Sense: 5′-<u>CTTC</u> GCAAGCCGAAACGAAGCTTC-3′

5’ Anti-sense: 5’-<u>AAAC</u> CAAGCTTCSTTTCGGCTTGC-3’

3’ Sense: 5′-<u>CTTC</u> GTCCGTCCGGAAGCGATAGA-3′

3’ Anti-sense: 5’ <u>AAAC</u> TCTATCGCTTCCGGACGGAC-3’

*Donor plasmid primers:*

5′ Homology arm (AarI sites):

Forward: 5′-tgcaCACCTGCgatcctac ATCCCAACTGAGCACAATGAG -3′

Reverse: 5′-attcCACCTGCtgaatcgc TTCTGGCACGCTTCGCTG -3′, 3′

Homology arm (SapI sites):

Forward: 5′-cgagGCTCTTCagac ATCGCTTCCGGACGGAC -3′

Reverse: 5′-tagtGCTCTTCatat TGATAGTGGTAGCGATACAC -3′

gRNA vectors and donor plasmids were sequence verified and co-injected into vasa-Cas9 embryos (BDSC#51324). Injections were performed by BestGene Inc. (California, USA). Adult survivors were crossed to balancer stocks, and F1 progeny were screened for dsRed fluorescence in the eyes using a fluorescent stereomicroscope. Positive individuals were selected as putative RalGPS deletion mutants. The marker cassette was removed by crossing to cyo,Crew (BDSC#1092), and the deletion was confirmed via sequencing. To generate endogenously tagged RalGPS-3xHA, CRISPR targeting was performed at the 3′ end of RalGPS using the guide sequence: 5’-cggcttattcaaaggacatt-3’. Editing and integration were performed by Rainbow Transgenic Flies Inc. (California, USA).

### Cloning of RalGPS for Transgenic Expression

The full-length RalGPS open reading frame (ORF) was amplified from cDNA (Drosophila Genomic Research Center, Indiana, USA) using: Forward primer: 5′-CACC ATGATGCGATACTCGGAAATCTC-3′, Reverse primer: 5′-TTA TTCAAAGGACATTAGGTTGGTGG-3′. The ORF was cloned into the pENTR vector using TOPO cloning (Invitrogen), and recombined into the pUASattB-10xUAS destination vector with either no tag or a N-terminal EGFP tag. Sequence verified constructs were injected into the attP2 genomic landing site by BestGene Inc.

### Generation of the RalGPS 3′UTR Fluorescent Reporter Line

A fluorescent transgenic reporter line was generated to assess RalGPS 3′UTR regulation by cloning a destabilized GFP (dGFP) followed by the RalGPS 3′UTR under the control of the ubiquitin promoter. To construct the reporter, the pUASattB destination vector was modified by replacing the UAS with the ubiquitin promoter (amplified from pUWR, DGRC#1281), followed by a dGFP sequence amplified from pUAST-IVS-syn21-nlsGFP-PEST-2A-nlsRFP-p10 (kindly provided by Perrimon lab;(He et al. 2019)). This generated a Gateway-compatible vector: pattB-Ubi-dGFP-attR-ccdb. The RalGPS 3′UTR was amplified from w^1118^ genomic DNA and cloned into a pENTR vector using TOPO cloning (Invitrogen). The entry clone was recombined into the pattB-Ubi-dGFP-attR-ccdb vector via Gateway cloning. Sequence-verified final constructs were injected into the attP40 genomic landing site by BestGene Inc. (California, USA).

### Immunohistochemistry and Western Blot

Wandering third instar Drosophila larvae were dissected in cold HL3 buffer and fixed in Bouin’s reagent for 5 min. Preparations were washed in PBST (PBS containing 0.1% Triton X-100), incubated with primary antibodies overnight at 4°C, and with secondary antibodies for 2 h at room temperature. Washes (3 x 15 min) were performed with PBST between incubations. Primary antibodies used were: mouse anti-Dlg1 (DSHB 4F3, 1:500), mouse anti-Synapsin (DSHB 3C11, 1:100), and rabbit anti-GFP (Invitrogen A-11120, 1:1000). Secondary antibodies included Alexa Fluor 488-or 568-conjugated antibodies (Invitrogen, 1:500) and HRP-conjugated antibodies (Alexa 594 or 647, 1:500). Images were acquired on Nikon A1R confocal and Nikon 90i fluorescence microscopes, processed and analyzed using Fiji (ImageJ) and Adobe Photoshop.

For Western blotting, dissected larval brains were lysed in IP lysis buffer (Invitrogen) supplemented with protease inhibitors (Roche). Lysates were denatured at 95°C for 10 min in 2 x Laemmli sample buffer (Bio-Rad), separated on 4–15% TGX gels (Bio-Rad), and transferred to PVDF membranes (Invitrogen). Membranes were blocked in 5% skim milk in TBST (TBS with 0.1% Tween-20) and incubated overnight at 4°C with rabbit anti-GFP (Invitrogen A-11120, 1:1000), rabbit anti-HA (Abcam ab9110, 1:1000), or mouse anti-Tubulin (DSHB E7, 1:250). After three 5-min TBST washes, membranes were incubated with HRP-conjugated secondary antibodies (Cell Signaling Technology, 1:3000) for 2 h at room temperature. Signals were detected using ECL (SuperSignal West Pico, Thermo Scientific) and developed on X-ray film (Fujifilm). Band intensities were quantified using Fiji and normalized to tubulin controls.

### RNA Seq : Sample Preparation, sequencing and data analysis

Wandering third instar Drosophila larvae were dissected in cold HL3 buffer. Larval pelts (excluding gut and fat body) were collected from control and miR-219 null genotypes, with 15 pelts pooled per biological replicate. Total RNA was extracted using the RNeasy Mini Kit (Qiagen) following the manufacturer’s instructions. RNA quality and concentration were assessed using the Agilent 2100 Bioanalyzer (Agilent Technologies), and samples with an RNA Integrity Number (RIN) ≥ 8 were used for library preparation. Libraries were prepared using poly(A) selection, and sequencing was performed by Novogene, targeting a minimum depth of 20 million reads per sample. Reads were aligned to the Drosophila melanogaster reference genome (r6.41, FlyBase) using STAR (Dobin et al. 2012). Resulting BAM files were indexed and quality-checked using SAMtools, followed by further processing with Picard v2.8.0. Read quantification was performed with featureCounts(Liao et al. 2014). Differential gene expression (DGE) analysis was conducted using the DESeq2 package in R (Love et al. 2014). Gene Ontology (GO) enrichment analysis was carried out using clusterProfiler (Yu et al. 2012), with annotations from org.Dm.eg.db (Carlson, 2019). Redundant GO terms were removed using the simplify function in clusterProfiler to enhance interpretability.

### AGO1 Immunoprecipitation and miRNA Analysis

Wandering third instar *Drosophila* larvae were dissected and subjected to high K⁺ (90 mM) depolarization as described above. Larval pelts (n = 20 per condition) were collected in lysis buffer (30 mM HEPES-KOH pH 7.4, 100 mM potassium acetate, 2 mM magnesium acetate, 5% glycerol, 0.5% IGEPAL CA-630, 1 mM EGTA, 1 mM DTT) supplemented with RNaseOUT (Invitrogen) and protease inhibitor cocktail (Roche). Samples were homogenized using a motorized pestle and incubated on ice for 10 min. Lysates were clarified by centrifugation at 15,000g for 10 min at 4°C, and supernatants were transferred to fresh tubes. For AGO1 immunoprecipitation, 3 µg of anti-AGO1 Fab fragments (kindly provided by H. Lipshitz, University of Toronto; generated as described in (Na et al. 2016)) were added to the lysates and incubated for 2 h at 4°C. Protein A/G magnetic beads (25 µL; MagnaChip) were added and incubated for 1 h at 4°C. Beads were washed 3 x in lysis buffer and transferred into TRIzol (Invitrogen) for RNA extraction using the miRNeasy Kit (Qiagen) according to the manufacturer’s instructions. A fraction (10%) of the beads was retained for validation of AGO1 immunoprecipitation by Western blot using mouse anti-AGO1 antibody (Miyoshi et al. 2005). Quantification of miRNAs from total RNA and AGO1 IP RNA was performed using TaqMan MicroRNA Assays (Thermo Fisher Scientific) on a QuantStudio 7 Pro (Applied Biosystems). 10ng of RNA was reverse transcribed using miRNA specific RT primers, which was further amplified by RT-PCR using universal Taqman master mix and miRNA specific primers. The relative abundance of miRNAs was calculated using the ΔΔCt method. The following probes were used: dme-miR-219-5p (#000522), dme-miR-34-5p (#464452_mat), dme-let-7-5p (#000332), and U14 snRNA (#001750) as internal control.

### mRNA expression analysis by quantitative PCR (qPCR)

Total RNA was extracted from dissected larval pelts using the RNeasy Mini Kit (Qiagen) according to the manufacturer’s instructions. Complementary DNA (cDNA) was synthesized from 25 ng of total RNA using the iScript cDNA Synthesis Kit (Bio-Rad). Quantitative PCR (qPCR) was performed on a QuantStudio 7 Pro RT-PCR System (Applied Biosystems) using iTaq™ SYBR® Green Supermix with ROX (Bio-Rad). Relative mRNA expression levels were calculated using the comparative Ct (ΔΔCt) method, with RpL32 serving as the internal normalization control.

Primer sequences used in this study are listed below:

### Statistical analysis

All statistical analyses were performed using GraphPad Prism version 10 (GraphPad Software, San Diego, CA). Appropriate statistical tests, including unpaired two-tailed Student’s t-test, one-way ANOVA, or two-way ANOVA, were applied as indicated in the figure legends. When necessary, post hoc corrections were applied to account for multiple comparisons. Data are presented as mean ± SEM. Statistical significance was defined as follows: **** P ≤ 0.0001, *** P ≤ 0.001, ** P ≤ 0.01, * P ≤ 0.05, and ns (P > 0.05; not significant).

---

**Figure S1:**
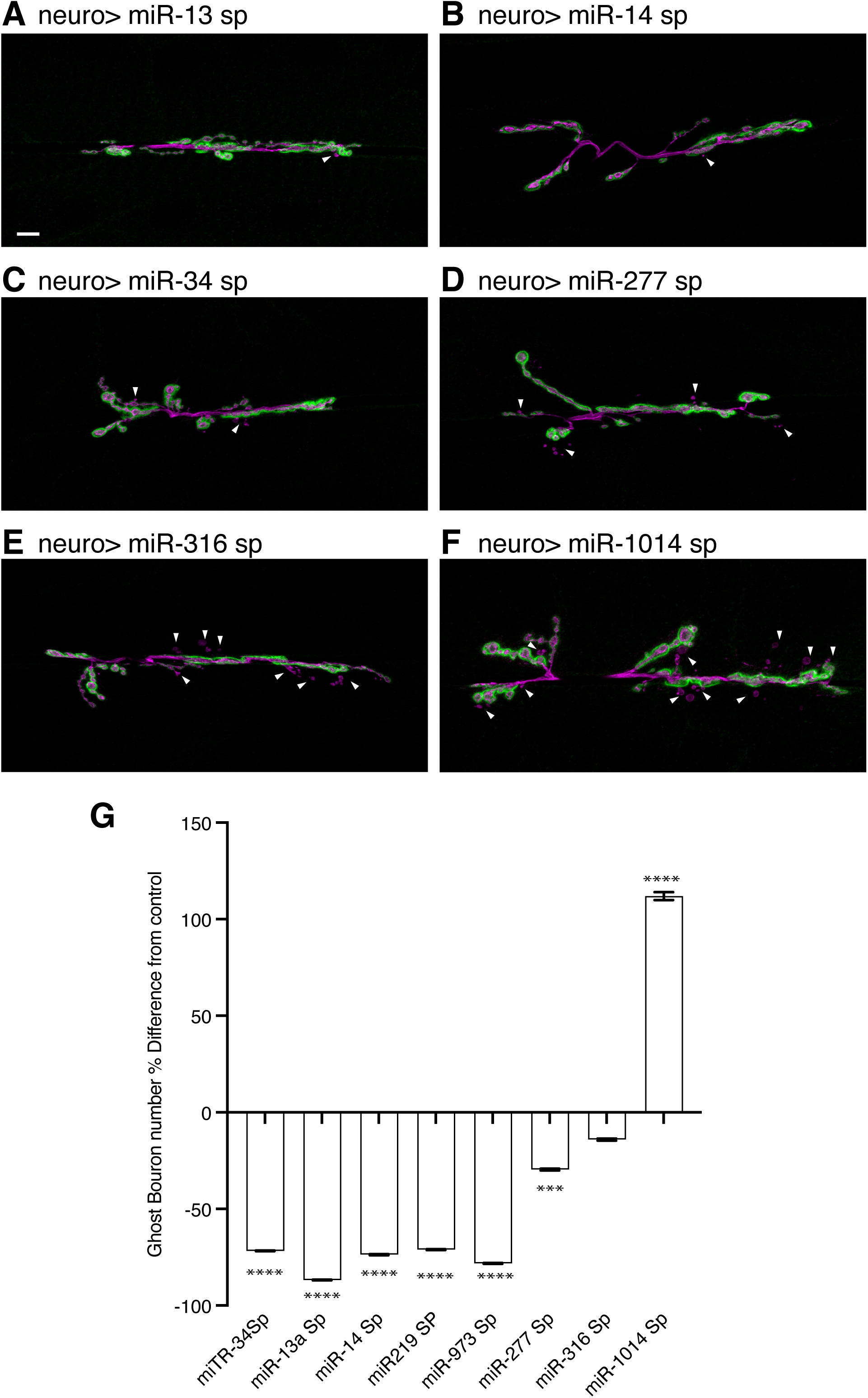
miRNA-dependent regulation of activity-induced ghost bouton formation. **A–F**. Confocal images of NMJ arbors on muscles 6 /7 after high K⁺ stimulation, stained for presynaptic membrane (HRP, magenta) and postsynaptic cytomatrix (Dlg, green). Arrowheads mark ghost boutons. miRNA sponges were expressed in motor neurons (OK371-GAL4/UAS-miR-sponge; UAS-miR-sponge/+). **A**. miR-13-5p **B**. miR-14-5p **C**. miR-34-5p **D**. miR-277-5p **E**. miR-316-5p **F**. miR-1014-5p. Scale bars: 10 μm. **G**. Quantification of ghost bouton induction (% difference from control across genotypes). Mean ± SEM, One-way ANOVA and Dunnett’s multiple comparisons. Statistical significance: ****p ≤ 0.0001***p ≤ 0.001.

**Figure S2:**
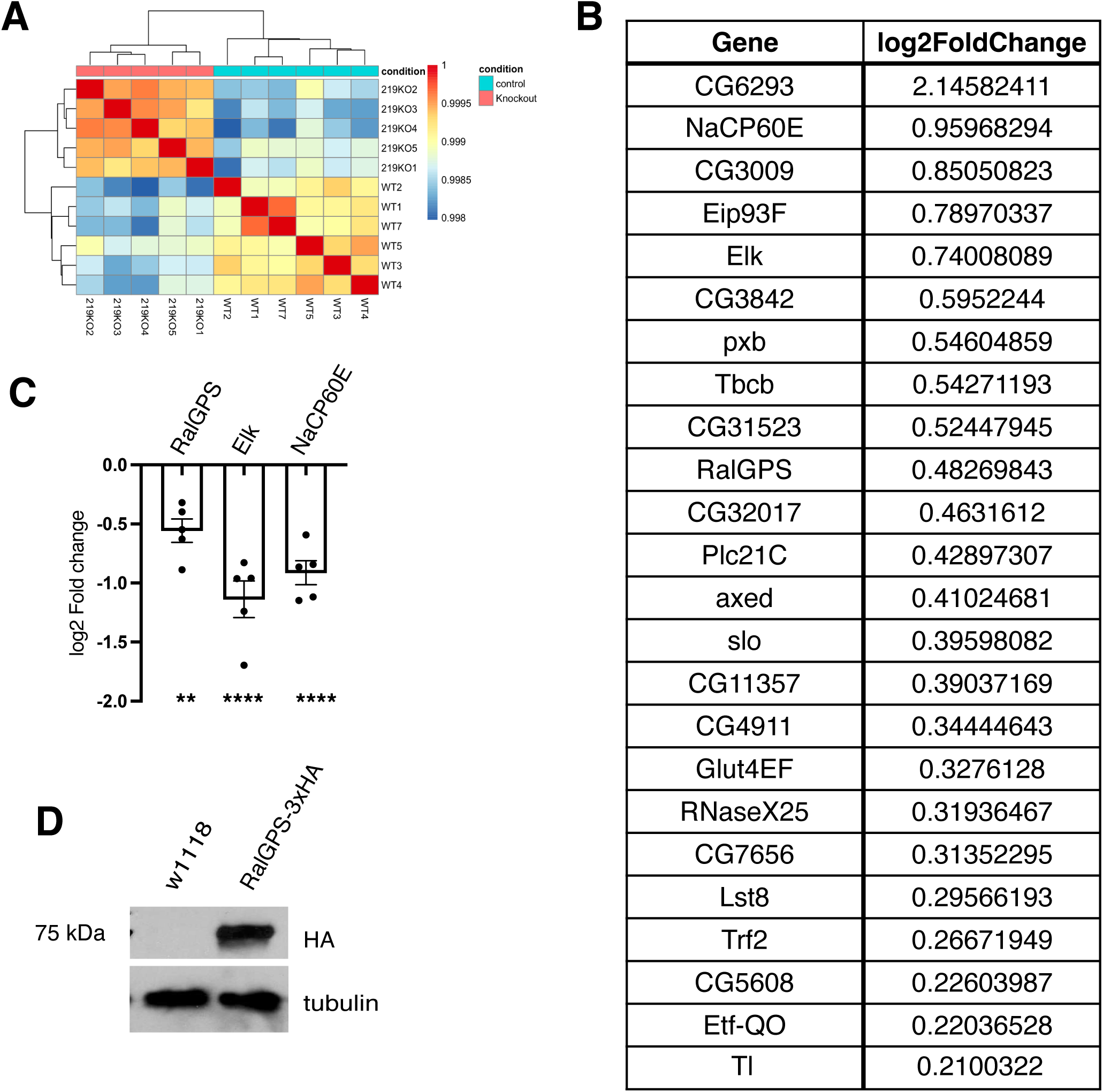
Transcriptome and molecular analysis of miR-219 and its target RalGPS. **A**. Hierarchical clustering of RNA-seq data from control and miR-219 null larvae. **B**. List of upregulated genes in miR-219 nulls containing predicted miR-219-5p MREs. **C**. qPCR showing downregulation of genes with conserved miR-219-5p MREs following high K+ treatment. Mean ± SEM, One-way ANOVA and Dunnett’s multiple comparisons. Statistical significance: ****p ≤ 0.0001, **p ≤ 0.01. **D**. Western blot showing RalGPS expression in brains from endogenously HA-tagged RalGPS flies.

**Figure S3:**
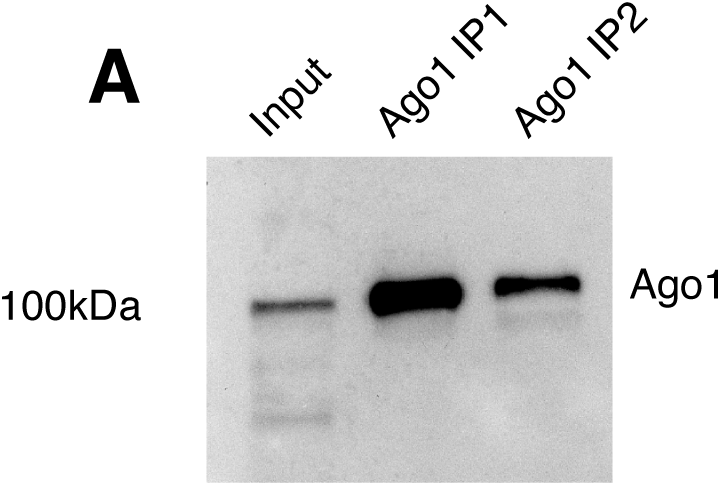
Validation of Ago1 immunoprecipitation. A. Western blot confirming Ago1 immunoprecipitation using two independent Ago1-specific Fabs, detected with mouse anti-Ago1 antibody.

**Figure S4:**
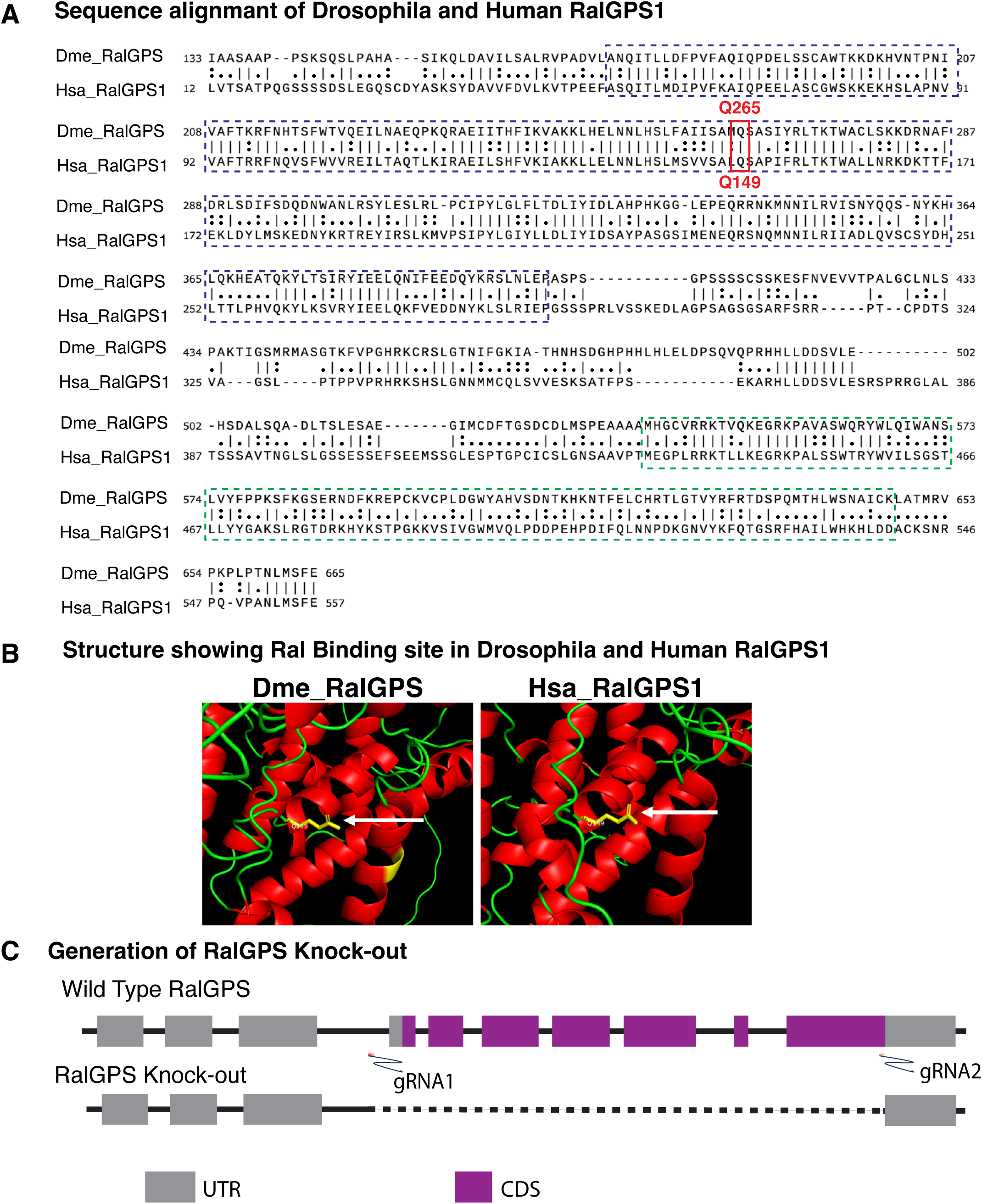
CRISPR-mediated generation of RalGPS null and conserved domain architecture between Drosophila and Human RalGPS. **A.** ClustalW sequence alignment of Drosophila RalGPS and human RalGPS1 highlighting the conserved Cdc25 domain (blue box) and PH domain (green box). The RalA-specific interaction site (glutamine residue) is indicated in red. **B.** Domain organization of Drosophila RalGPS and human RalGPS1 showing the conserved RalA-specific interaction residue (white arrow showing Gln265 in Drosophila and Gln149 in human). **C.** Schematic of the CRISPR/Cas9-mediated genomic deletion strategy used to generate the RalGPS null allele.

